# Brain State-Dependent Modulation of Thalamic Visual Processing by Cortico-thalamic Feedback

**DOI:** 10.1101/2021.10.04.463017

**Authors:** Kimberly Reinhold, Arbora Resulaj, Massimo Scanziani

**Affiliations:** Neurosciences Graduate Program, University of California San Diego, La Jolla, California, USA; Center for Neural Circuits and Behavior, Neurobiology Section and Department of Neuroscience, University of California San Diego, La Jolla, California, USA; Department of Physiology, University of California San Francisco, San Francisco, California, USA; Howard Hughes Medical Institute, University of California San Francisco, San Francisco, California, USA

**Author notes:** These authors contributed equally (Kimberly Reinhold, Arbora Resulaj).

## Abstract

The behavioral state of a mammal impacts how the brain responds to visual stimuli as early as in the dorsolateral geniculate nucleus of the thalamus (dLGN), the primary relay of visual information to the cortex. A clear example of this is the markedly stronger response of dLGN neurons to higher temporal frequencies of the visual stimulus in alert as compared to quiescent animals. The dLGN receives strong feedback from the visual cortex, yet whether this feedback contributes to these state-dependent responses to visual stimuli is poorly understood. Here we show that in mice, silencing cortico-thalamic feedback abolishes state-dependent differences in the response of dLGN neurons to visual stimuli. This holds true for dLGN responses to both temporal and spatial features of the visual stimulus. These results reveal that the state-dependent shift of the response to visual stimuli in an early stage of visual processing depends on cortico-thalamic feedback.

## Introduction

While awake, animals shift between brain states: the alert state, when animals engage with sensorimotor behaviors, and a quiescent state (Bezdudnaya et al., 2006; O’Connor et al., 2002; Steriade, 1997). These shifts affect visual processing as early as the dorsolateral geniculate nucleus of the thalamus (dLGN). For example, in the alert state, dLGN neurons respond better to visual stimuli that change rapidly in time, i.e., containing high temporal frequencies, as compared to the quiescent state (Bezdudnaya et al., 2006; Aydin et al., 2018). Numerous mechanisms have been proposed for these state-dependent shifts in visual processing (McCormick, 1992; Castro-Alamancos, 2004; Hirata et al., 2016; Briggs et al., 2008; Fanselow et al., 2001, Eyding et al., 2003, Ruiz et al., 2006, Molnár et al., 2021; de Labra et al., 2007; Huguenard and McCormick, 2007), yet the contribution of the cortical feedback projections to the dLGN has not been established.

Visual cortex sends massive feedback projections to the dLGN via specific excitatory neurons located in layer 6 (Crandall et al., 2015; Destexhe, 2000). In fact, these projections outnumber the retinal input by an order of magnitude (Van Horn et al., 2000). Because visual cortex undergoes dramatic shifts in activity between the alert and quiescent states (Niell and Stryker, 2010; Lee et al., 2014), visual cortex, through its feedback to the thalamus, may well impact the visual responses of the dLGN in a state-dependent manner.

While many studies have addressed the role of cortico-thalamic feedback on visual responses in the dLGN (Andolina et al., 2007; Crandall et al., 2015; Denman and Contreras, 2015; McAlonan et al., 2008; Mease et al., 2014; O’Connor et al., 2002; McAlonan et al., 2006; Denman et al., 2015; Kirchgessner et al., 2020; Born et al., 2021), none of these studies tested whether cortico-thalamic feedback contributes to the differences in the visual response properties of the thalamus depending on the state of the animal.

Here we use optogenetic approaches to rapidly and reversibly silence either visual cortex or the layer 6 cortico-thalamic neurons, while simultaneously recording from the dLGN and the hippocampus to record visual responses and to monitor brain state, respectively. Silencing feedback projections removed brain state-dependent differences in the temporal and spatial response properties of dLGN neurons, without affecting brain state. Therefore, feedback projections from visual cortex to the thalamus contribute to the state-dependent shifts in how the external world is represented even at this very early stage of sensory processing.

## Results

### Defining alert and quiescent brain states

We measured global brain states in awake mice that were head-fixed yet free to spontaneously run or rest on a circular treadmill (Methods) (Reinhold et al., 2015). We defined brain state by the presence or absence of hippocampal theta oscillations in the local field potential (LFP) recorded just ventral of the hippocampus, consistent with previous descriptions (Bezdudnaya et al., 2006). The power of the theta frequency band (5-7 Hz) was bimodally distributed and inversely correlated with the power of low-frequency oscillations (0.5-6 Hz) simultaneously recorded in visual cortex (Figure 1 A,B,C). We refer to periods of high power in the hippocampal theta frequency band as “alert state”, a brain state that has been linked to running, enhanced alertness and high sensorimotor engagement (Buzsáki, 2002; Green and Arduini, 1954). Conversely, periods of low power in the hippocampal theta frequency band, generally associated with reduced alertness to external sensory stimulation (Bereshpolova et al., 2011; Bezdudnaya et al., 2006; Kandel and Buzsáki, 1997; Swadlow and Gusev, 2001), are here referred to as “quiescent state”. During the alert state, the mice were mostly but not always running (Figure 1 B; in 86% of alert trials, the mouse was running; also see Figure Supplement 1 and Methods for a more detailed description of these two brain states), while during the quiescent state the mice were almost never running (the mice ran in <4% of trials classified as quiescent). The alert and quiescent states alternated every 9.5 seconds on average (9.5±13.0 s, mean ± std. dev.) and lasted on average 7.6 seconds and 11.7 seconds, respectively (alert: 7.6±6.4 s, mean ± std. dev., quiescent: 11.7±18.3 s, mean ± std. dev.).

**Figure 1.**
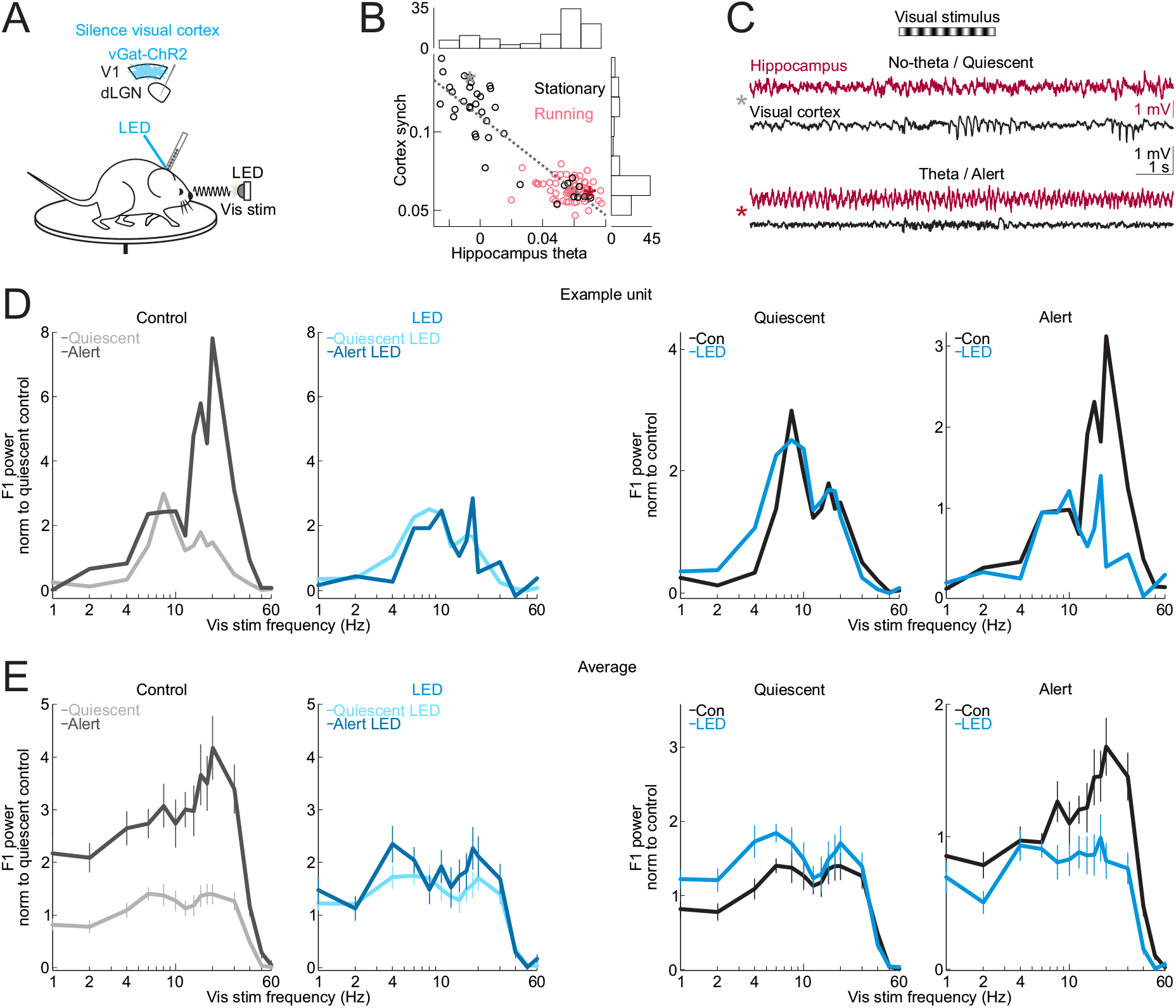
Silencing the visual cortex removed brain state-specific features of the visual responses of dLGN neurons. **(A)** Awake head-fixed mouse free to run or rest on a circular treadmill during electrophysiology recording, presentation of a full-field, flickering visual stimulus, and silencing of visual cortex by LED illumination of ChR2-expressing inhibitory cortical interneurons (vGat-ChR2). **(B)** Each dot is average across one 14.5 second long trial. All trials from example recording session in 1 mouse (Figure Supplement 1 for summary across 4 mice). Hippocampus theta (hippocampal LFP power at 5-7 Hz minus power at 2-4 Hz) vs. cortex synchronization (V1 LFP power at 0.5-6 Hz minus power at 30-80 Hz) (Methods). Black (non-running) vs. pink (running) indicates the animal’s motor behavior on each trial. Histograms show the distributions of hippocampus theta and cortex synchronization along each axis. **(C)** Example single trials of simultaneously recorded hippocampal LFP (dark red) and visual cortex LFP (black) before, during and after presentation of drifting grating visual stimulus (black and white stripes above traces). Top 2 traces: trial during the quiescent brain state (gray star in scatter in (B)). Bottom 2 traces: trial during the alert brain state (dark red star in (B)). **(D)** Example frequency response of a single unit recorded in dLGN. Experimental set-up as described in (A). Visual stimulus: full-field flicker at temporal frequencies from 1 to 60 Hz. **(E)** Effect of silencing cortical activity on mean and s.e.m. power of dLGN F1 frequency response across single units (n=254 units from 5 mice) during presentation of full-field flicker at temporal frequencies on X axis. F1 power was baseline-subtracted and normalized (Methods) to response across all frequencies.

### Contribution of visual cortex to brain state dependence of thalamic responses to visual stimuli

To determine how brain state impacts the response of dLGN neurons to temporal features of the visual stimulus, we recorded from these neurons using multi-site extracellular probes. The dLGN recording sites were labeled with a fluorescent dye and validated *ex vivo* (Methods). To hold the spatial structure of the visual stimulus constant while varying the temporal frequency, we presented a visual stimulus called full-field flicker, the sinusoidal modulation of full-field luminance at temporal frequencies between 1 and 60 Hz (Methods), to the eye contralateral to the recorded hemisphere.

The firing of isolated putative dLGN relay neurons (n=254 units from 5 mice) was modulated by the fundamental temporal frequency (F1) of the visual stimulus. For example, in response to a visual stimulus flickering at 4 Hz, the firing rate of dLGN neurons was modulated at 4 Hz (44% of the visually modulated spikes were F1-modulated, 30% were F2-modulated, <26% were modulated at other frequencies, Figure Supplement 2). We defined the F1 response of a neuron as the amplitude of the F1 modulation of its firing rate in response to a range of stimulus frequencies. The F1 response of dLGN neurons was markedly different between the alert and quiescent states, consistent with results in other species (Bezdudnaya et al., 2006) and previous work (Aydin et al., 2018). Not only was the F1 response more pronounced during the alert state, but it also peaked at higher temporal frequencies (Figure 1 D and E, left-most panels). That is, in the alert state, the activity of dLGN neurons was more strongly driven by the temporal properties of the visual stimulus and followed higher frequencies, as compared to the quiescent state. Thus, the temporal response of mouse dLGN neurons to visual stimuli depends on brain state.

Does primary visual cortex (V1) contribute to the dependence of dLGN neurons’ responses to brain state? To address this question, we optogenetically silenced V1 using the vGat-ChR2 mouse line, in which the opsin is selectively expressed in GABAegic neurons that inhibit V1 projection neurons (Olsen et al., 2012; Methods). V1 was silenced (Figure Supplement 3) for a brief period of time (3 second window beginning 0.5 seconds before the onset of the visual stimulus and continuing throughout the visual stimulus) during either the alert or quiescent state while presenting visual stimuli at various temporal frequencies. V1 illumination did not change brain state (Figure Supplement 4), as measured by the theta power of the LFP of hippocampal recordings, nor did it impact running behavior (switch from stationary to running or running to stationary in 20% of trials without V1 silencing and 16% of trials with V1 silencing, n.s., p-val=0.22 from Wilcoxon sign-rank across sessions) consistent with previous studies (Resulaj et al., 2018). V1 illumination also did not affect the power of the LFP of nearby primary somatosensory cortex (Figure Supplement 4 C). Silencing V1 reduced spontaneous, non-visually evoked dLGN activity in both the quiescent and alert states (quiescent state: (mean ± s.e.m) from 5.1±0.24 Hz to 3.9±0.21 Hz, alert state: from 7.1±0.32 Hz to 4.9±0.24 Hz, n = 377 units, 6 mice; Figure Supplement 5 A) and increased burst activity, also in both brain states (Figure Supplement 5 B,C,D,E). Strikingly, silencing V1 eliminated brain state differences in the visually evoked F1 response of dLGN neurons (Figure 1 D,E and Figure Supplement 2). V1 silencing strongly reduced the dLGN response to high frequencies (10 – 60 Hz) in the alert state while increasing the response to low frequencies (1 – 8 Hz) in the quiescent state (alert state: 41% decrease in the response to high stimulus frequencies, quiescent state: 39% increase in the response to low stimulus frequencies, n=254 units from 5 mice; Figure Supplement 2 B,F,G for statistics). As a consequence, the F1 response in the alert and quiescent states became almost indistinguishable (Figure 1 E and Figure Supplement 6). These results show that silencing V1, by reducing the F1 response to high frequencies in the alert state and enhancing the response to low frequencies in the quiescent state, effectively eliminates the visual response features that are unique to each brain state and leads to a response function that is intermediate between the quiescent and alert brain states. Thus, the state-dependent differences in the F1 response of the dLGN depend on V1.

### Contribution of the cortico-thalamic feedback projection

To determine whether V1 impacts the F1 response of the dLGN via direct cortico-thalamic projections, we took advantage of a mouse line (Ntsr1-Cre) that selectively labels layer 6 cortical neurons that project back to the dLGN (Bortone et al., 2014) and conditionally expressed either the inhibitory opsin GtACR2 or the inhibitory opsin ArchT in this population of cortico-thalamic neurons (Methods) to suppress them. Consistent with V1 silencing, photo-suppression of these layer 6 cortico-thalamic neurons decreased spontaneous dLGN activity in both the quiescent and alert brain states, albeit to a lesser extent, likely because of the incomplete suppression of cortico-thalamic neurons (quiescent state (mean ± s.e.m.): spontaneous dLGN activity from 3.9±0.25 Hz to 3.3±0.23 Hz, alert state: from 5.4±0.31 Hz to 4.9±0.30 Hz, n = 231 units, n = 7 mice, quiescent state: p < 10^-6^, alert state: p < 10^-6^ from paired Wilcoxon sign-rank test across units). Importantly, however, for dLGN neurons showing a significant reduction of baseline activity upon layer 6 silencing, the F1 response to low temporal frequencies in the quiescent brain state was enhanced, while the F1 response to high temporal frequencies in the alert brain state was suppressed (alert state: 45% decrease to high stimulus frequencies, quiescent state: 143% increase to low stimulus frequencies, n=43 units from 4 mice; Figure 2; Figure Supplement 7; Figure Supplement 8). These data indicate that the state-dependent differences in the F1 response of the dLGN are mediated by V1 through cortico-thalamic layer 6 neurons.

**Figure 2.**
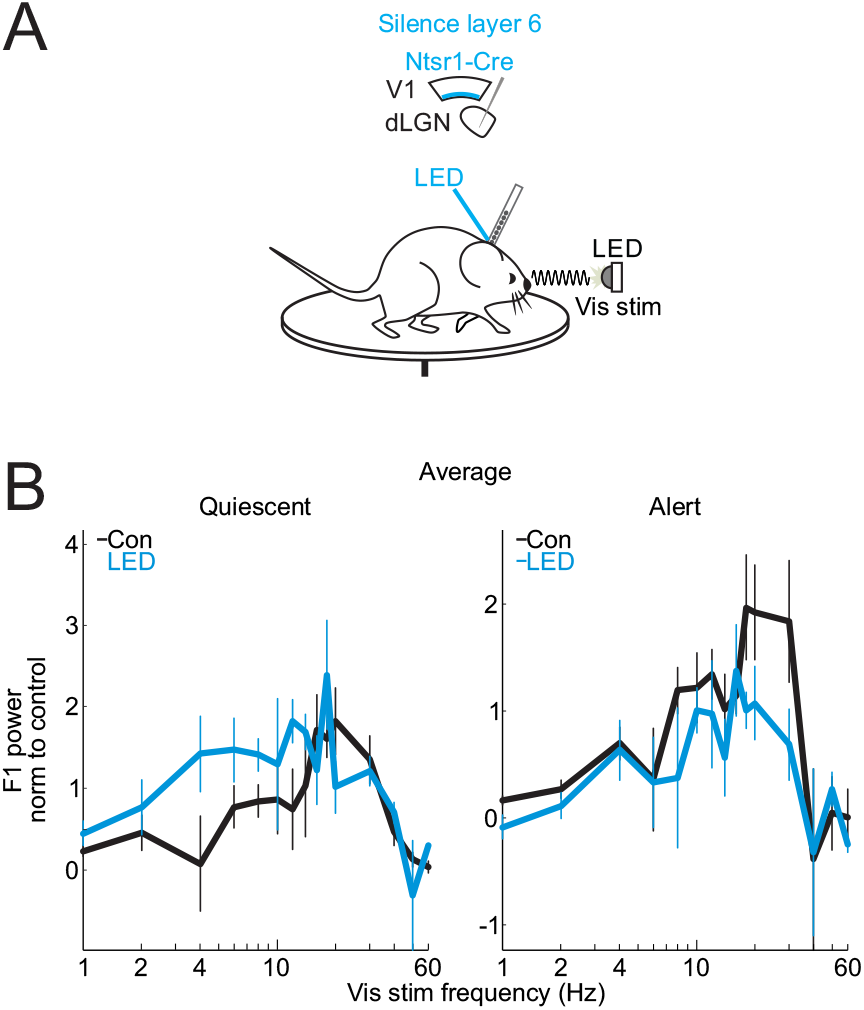
Suppressing cortico-thalamic feedback from visual cortex layer 6 reduces brain state-specific features of the visual responses of dLGN neurons. **(A)** Awake head-fixed mouse free to run or rest on a circular treadmill during electrophysiology recording, presentation of a full-field, flickering visual stimulus, and suppression of visual cortex layer 6 Ntsr1+ cortico-thalamic neurons expressing the inhibitory opsin GtACR2. **(B)** Effect of suppressing layer 6 cortico-thalamic neurons on mean and s.e.m. power of dLGN F1 frequency response across single units (n=43 units from 4 mice) during presentation of full-field flicker at temporal frequencies on X axis. F1 power was baseline-subtracted and normalized (Methods) to response across all frequencies. Figure includes only dLGN units with at least a 20% suppression of spontaneous activity at LED onset (Methods). See Figure Supplement 7 D for summary across all units.

### Impact of visual cortex on visual spatial frequency tuning in dLGN

We next investigated whether brain state also affects the response of the dLGN to varying spatial frequencies of the visual stimulus, and if this also depends on cortico-thalamic feedback. We used drifting gratings presented on a computer monitor to the eye contralateral to the recorded hemisphere (Figure 3 A) and varied the spatial frequency of the gratings (0.02-0.08 cycles per degree) while holding the temporal frequency constant (3 Hz). The spatial frequency response of dLGN neurons was larger in the alert state compared to the quiescent state for frequencies around 0.03 cycles per degree (cpd) (at 0.03 cpd: alert state response was 79% larger than quiescent state response) indicating that brain state also affects the spatial frequency response of these neurons. Strikingly, silencing visual cortex, again, abolished this difference (Figure 3 B). This was mainly due to an overall increase in the responses recorded in the quiescent state across all spatial frequencies (silencing V1 in the quiescent state increased response by 174% at 0.02 cpd, 167% at 0.03 cpd, 118% at 0.04 cpd, and 70% at 0.08 cpd; silencing V1 in the alert state increased response by 136% at 0.02 cpd, 24% at 0.03 cpd, 57% at 0.04 cpd, and 330% at 0.08 cpd; see Figure Supplement 9 for full distributions of units, preferred vs. non-preferred tuning and statistics). Thus, silencing visual cortex enhances the spatial frequency response of the dLGN in the quiescent state to match that of the alert state. Similar to silencing visual cortex generally, suppressing specifically layer 6 cortico-thalamic neurons expressing ArchT increased the spatial frequency responses of dLGN neurons preferentially in the quiescent state (Figure 4, Figure Supplement 10).

**Figure 3.**
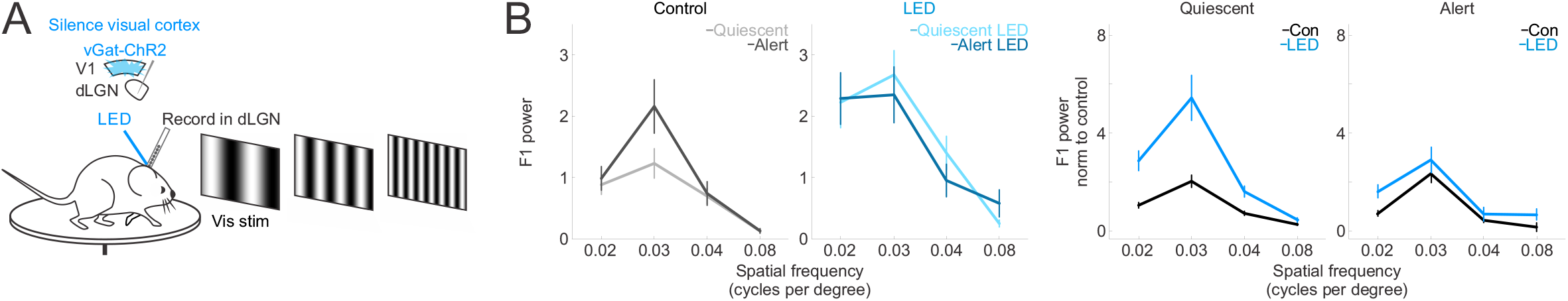
Silencing the visual cortex removes the brain state-specific differences in the spatial frequency response of dLGN neurons. **(A)** Awake head-fixed mouse free to run or rest on a circular treadmill during electrophysiology recording, visual stimulus presentation of drifting gratings with various spatial frequencies, and silencing of visual cortex by LED illumination of ChR2-expressing inhibitory cortical interneurons. **(B)** Effect of silencing cortical activity on mean and s.e.m. of dLGN F1 response across dLGN single units (n=238 units from 5 mice) at various spatial frequencies of the drifting grating. Left two plots: raw F1 power in units of 1e5. Right two plots: F1 power was baseline-subtracted and normalized (Methods) to response across all frequencies as in Figure 1. See Figure Supplement 9 for distributions of units.

**Figure 4.**
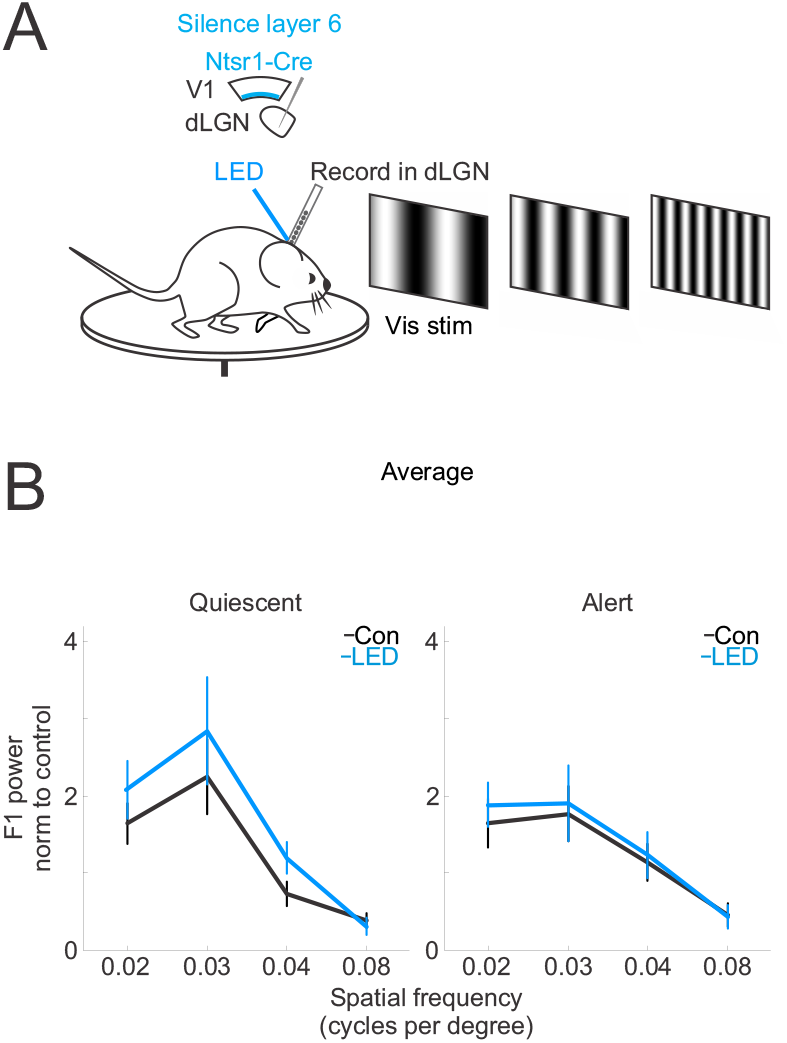
Suppressing cortico-thalamic feedback from visual cortex layer 6 reduces the brain state-specific differences in the spatial frequency response of dLGN neurons. **(A)** Awake head-fixed mouse free to run or rest on a circular treadmill during electrophysiology recording, visual stimulus presentation of drifting gratings with various spatial frequencies, and suppression of ArchT-expressing layer 6 cortico-thalamic neurons. **(B)** Effect of suppression of layer 6 cortico-thalamic neurons on mean and s.e.m. of dLGN F1 response across dLGN single units (n=195 units from 5 mice) at various spatial frequencies of the drifting grating. F1 power was baseline-subtracted and normalized (Methods) to response across all frequencies as in Figure 1. See Figure Supplement 10 for distributions of units.

## Discussion

The transmission of visual information from the retina to the cortex relies on an intermediary thalamic nucleus, the dLGN. The response properties of the dLGN to visual stimuli are highly dependent on the state of alertness of the animal (Bezdudnaya et al., 2006; Aydin et al., 2018; Williamson, 2015). Our results indicate that a major component of this state-dependence of dLGN responses requires the activity of visual cortex and, specifically, the feedback projections from cortex to thalamus. Indeed, when silencing visual cortex, the response properties of dLGN to visual stimuli are similar across states of alertness. In the temporal domain, this equalization of responses across states results from a reduction of the F1 response to high frequencies in the alert state and an enhancement of the F1 response to low frequencies in the quiescent state. As a consequence, upon cortical silencing, the temporal response properties of the dLGN are intermediate between the quiescent and alert brain states, irrespective of the actual brain state of the animal. In the spatial domain, the equalization of the responses between alert and quiescent states results from an enhancement of the response, in particular, to middle frequencies in the quiescent state.

Although silencing visual cortex decreases and enhances F1 responses in the alert and quiescent states, respectively, it reduces spontaneous activity in the dLGN irrespective of brain state. Thus, in the absence of visual stimulation, the net effect of visual cortex onto dLGN neurons is excitatory irrespective of brain state (Figure Supplement 5 A). In the presence of visual stimulation, however, the effect of cortical silencing on dLGN responses depends on the features of the visual stimulus. Therefore, cortical feedback is unlikely to simply scale the amplitude of thalamic responses (i.e., gain scaling or divisive inhibition). Silencing visual cortex increased the bursting frequency of dLGN neurons regardless of brain state (Figure Supplement 5 E). This bursting of dLGN neurons results from a combination of intrinsic and circuit properties that, together, may make dLGN neurons more likely to follow low frequency as compared to high frequency retinal input (Steriade, 1996; McCormick, 1990; McCormick and Huguenard, 1992). This may explain the shifts from preferring higher to lower temporal frequencies upon cortical silencing. Alternatively, or in addition, in the alert state, cortex is responsive to the temporal properties of the stimulus, while in the quiescent state, cortex is dominated by its own internal activity (Steriade, 1997; Vinck et al., 2015; Nestvogel, 2021; Clancy, 2019; Clayton, 2021). As a consequence, in the alert state, the cortical feedback may enhance visual responses in the dLGN, while in the quiescent state, the cortical feedback may actively disrupt dLGN neurons’ visual responses by adding a pattern of activity that is not matched to the visual stimulus. Future studies will eventually provide support for or against this hypothesis.

Our identification of the contribution of cortical feedback brings us closer to understanding how the brain coordinates activity across cortex and thalamus to switch from an externally oriented brain state for sensory processing, in which the top-down feedback enhances the rapidly changing feed-forward sensory signal, to an internally oriented brain state, in which the top-down feedback suppresses the sensory response, possibly shifting the thalamus toward a representation of internal processes such as expectation, imagery or working memory.

## Acknowledgements

We thank M. Mukundan and J. Evora for help with mouse husbandry. This project was supported by the NIH R01EY025668, K99EY030415 (A.R.) and Howard Hughes Medical Institute.

## Author Contributions

K.R. and M.S. designed the experiments. K.R. and A.R. performed the experiments. K.R. analyzed the data. K.R., A.R. and M.S. wrote the manuscript.

## Experimental Procedures

### Animal protocols

All procedures were conducted in accordance with the National Institutes of Health guidelines and with the approval of the Committee on Animal Care at UCSD (protocol S02160M) and UCSF (protocol AN131913). Animals were housed on a reverse light cycle in cages of five mice or less. At the time of electrophysiology, all animals were older than 3.5 weeks. Both male and female animals were used in an approximately equal ratio.

### Mouse lines

vGat-ChR2 (Jackson Labs stock number: 014548), Ntsr1-Cre

### Viruses

AAV2/9.CAG.flex.Arch.GFP from the University of North Carolina viral vector core AAV2/1.CAG.flex.ChR2.GFP from the University of North Carolina viral vector core AAV2/9.hSyn1.Flex.GtACR2.FusionRed prepared at Janelia Research Campus (Titer used: 5*10^11)

### Animal surgery: awake recordings

We implanted a chronic head frame for awake electrophysiology recordings at least 5 days before the recording.

### Head frame surgery

Animals were anesthetized with 2% isoflurane. After removing the hair on the head and sterilizing the skin, the skin on the top part of the skull was removed. A bone scraper and a scalpel blade were used to clean and score the surface of the skull. A bone screw was inserted bilaterally into each skull plate at 0.75 mm anterior and 2 mm lateral of bregma. Then, a thin layer of Vetbond was applied to the skull followed by the application of dental cement mixed with black paint to affix the head frame to the skull. The black dental cement covered all areas of the skull except primary visual cortex to prevent light for optogenetic stimulation from accessing other cortical areas. This dental cement was also used to build up a recording well for containing artificial cerebrospinal fluid (ACSF) during in vivo electrophysiology. A single dose of 10% buprenorphine was administered, and we checked on the mice daily after the surgery.

### Electrophysiology in dLGN and V1

Before recording, we briefly anesthetized the mice to drill away the dental cement covering the skull, as previously described (Reinhold et al., 2015), to thin the skull over V1, and to make two small craniotomies (each about 50 microns in diameter) over V1 and over the dorsal lateral geniculate nucleus (dLGN). We thinned the skull over V1 until it was transparent when coated in ACSF. We targeted the V1 recording to approximately 3.75 mm posterior and 2.5 mm lateral of bregma, and we targeted the dLGN recording to approximately 1.8 mm posterior and 2 mm lateral of bregma, and 2.4 to 3 mm beneath pia, measuring visual responses to help target the recording sites in visual thalamus. For these recordings, we placed the V1 and dLGN electrodes in the brain before allowing the animal to wake up from anesthesia.

### Marking recording track in visual thalamus

The electrode placed in visual thalamus was coated with DiI to label the recording track, as previously described (Reinhold et al., 2015). Recording sites in thalamus were then verified post-mortem (see below).

### Recording the S1 local field potential (LFP)

We measured the LFP in primary somatosensory cortex (S1), with a 16-channel linear extracellular array from a site approximately 250 μm beneath the pia.

### Animal surgery: viral injections

#### Pup injections of ArchT into Ntsr1-Cre visual cortex

We injected AAV2/9.CAG.flex.Arch.GFP into the left visual cortex of neonatal mouse pups between postnatal days 0 and 2, as previously described (Reinhold et al., 2015). We did not record from these animals before post-natal day 30.

#### Adult injections of GtACR2 into Ntsr1-Cre visual cortex

We performed 3 craniotomies per mouse in the left hemisphere only. The craniotomies were located at (1) [0.4 mm anterior of lambdoid suture, 2.35 mm lateral of midline], (2) [0.4 mm anterior of lambdoid suture, 2.85 mm lateral of midline], and (3) [1.2 mm anterior of lambdoid suture, 2.6 mm lateral of midline]. Within each craniotomy we performed one injection of 150 nl at each of two depths: 700 um and 450 um. The injection speed was 30 nl/min. Recordings were done at 2 weeks post-injection because of virus toxicity observed at 3+ weeks.

### Post-mortem histology

Animals were not perfused, but brains were fixed in 4% paraformaldehyde (PFA) in phosphate-buffered saline (PBS) for 2 nights at 4 °C. Brains were then transferred to 30% sucrose for 2 days before sectioning on a freezing microtome. Sections were 50 μm thick. We then mounted the sections with a mounting medium containing DAPI. Sections were imaged using a fluorescent microscope.

### Verification of thalamic recording sites

Electrophysiology recording tracks targeted to thalamus were marked with DiI at the time of the recording (see above). Penetration of the dLGN was confirmed post-mortem. We discarded the data from mice in which we observed a recording track outside of dLGN.

### Habituating mice to recording set-up

Before recording, we placed each mouse on a circular treadmill in the head-fix set-up for 30-60 min each day for 3-5 days. Once the mouse appeared comfortable in the head-fix, we proceeded with recordings. Visual stimuli consisting of drifting gratings were presented during habituation sessions.

### Measuring running

We attached a rotary encoder to the circular treadmill to measure the animal’s running speed and direction.

### Extracellular electrophysiology

At each recording site in visual cortex or dLGN, we used either 1) a 16-channel linear NeuroNexus silicon probe (A series), with either 50 μm or 25 μm spacing between linearly arranged electrode sites, or 2) a 32-channel linear NeuroNexus silicon probe with 20 μm spacing between linearly arranged electrode sites. The 50 μm spacing in the 16-channel linear probe enabled simultaneous recordings from all V1 layers. We used the 25 μm spacing to specifically target deeper V1 layers. Recording electrodes were connected to a 10X or 20X pre-amplifying head-stage from AM Systems (Plexon adapter). The dLGN recording was then amplified 200X using the AM Systems 3500 amplifier. The V1 recording was amplified 500X using the AM Systems 3600 amplifier. Both extracellular voltage traces were filtered between 0.1 Hz and 10 kHz at the amplification step. A National Instruments Data Acquisition device was used to digitize the analog voltage signals, before these signals were acquired and displayed in Matlab (custom software by S. Olsen and K. Reinhold, as in (Reinhold et al., 2015)).

### Visual stimulation

Two types of visual stimulation were used:

#### Drifting gratings

Sinusoidal drifting gratings were oriented patterns of periodically varying luminance (i.e., black and white bars with graded transitions). Spatial frequency and temporal frequency (i.e., drift speed) were varied for the drifting gratings. Unless otherwise specified, figures show an average of responses across spatial frequencies 0.02, 0.03, 0.04, 0.06 and 0.08 cycles per degree and a temporal frequency of 3 Hz. For drifting gratings, we used an LCD computer monitor for display (gamma-corrected, mean luminance 50 cd/m^2, refresh rate 60 Hz) at a distance of 25 cm from the eye contralateral to the V1 recording site. Matlab’s Psychophysics Toolbox was used to display the drifting gratings. Unless otherwise specified, drifting grating contrast was 100%, and the mean luminance of the drifting grating was matched to the gray inter-stimulus blank screen.

#### Full-field flicker

By positioning a blue light-emitting diode (LED) in front of a lens (5×, 0.15 NA), we placed an approximately 1⁄2 inch homogeneous beam over the eye of the mouse contralateral to the recording sites. There was no spatial structure to this beam. We flickered this LED at varying temporal frequencies between 1 Hz and 60 Hz. Unless otherwise specified, the contrast of this flicker was 100%, varying between 0 mW and 25 mW (approx. 3.5 mW/cm^2). The luminance varied sinusoidally in time.

### Measuring spontaneous activity

We measured spontaneous activity during the presentation of an unchanging blank gray screen on the computer monitor (see above).

### Photo-activation of cortical inhibitory interneurons to silence V1 excitatory cells

A blue LED (either 455 nm or 473 nm) coupled to an optical fiber with a 1 mm diameter was positioned just above the visual cortex, ipsilateral to the dLGN recording. The skull was thinned above V1 and coated in ACSF on the day of the electrophysiology experiment (see “Animal surgery: awake recordings”), allowing the blue light from the LED (25 mW total power) to penetrate the cortex. Because the vGat-ChR2 transgenic mouse line expresses ChR2 in vGat+ inhibitory interneurons in the cortex, blue light in cortex activates these inhibitory interneurons. The illumination of V1 inhibited putative excitatory neurons in cortex (Figure Supplement 3 and as in (Reinhold et al., 2015)).

### Photo-inhibition of Ntsr1+ neurons in visual cortex

#### ArchT

Either an amber LED (595 nm, 20 mW) coupled to an optical fiber with a 1 mm diameter or an amber laser (550 nm, 25 mW) coupled to a fiber with a 0.2 mm diameter was positioned just above the visual cortex, ipsilateral to the dLGN recording. The skull was thinned over V1, as described above. Illumination of visual cortex led to suppression of neurons in V1 (Figure Supplement 8 B). We included data from experiments using both the amber LED and the amber laser.

#### GtACR2

We used a blue LED (25 mW) coupled to an optical fiber with a 1 mm diameter to illuminate the visual cortex (Figure Supplement 7 B).

### Blue light control

We used ambient blue light surrounding the mouse to mask sight of the blue LED used for optogenetic manipulation. Additionally, to test whether the blue light emitted by the optogenetic LED could affect the visual response directly (i.e., via the retina), in one mouse, we occluded the craniotomy with opaque Kwik-Cast during the recording, after verifying an effect of optogenetic cortical silencing on dLGN activity. The blue optogenetic LED remained positioned above the visual cortex but was unable to illuminate the cortex through the opaque Kwik-Cast. We tested whether turning on the LED in this case, in the absence of any optogenetic manipulation, had an effect on the visual response in dLGN. There was no effect (not shown). Similarly, when using the amber LED/laser to inhibit ArchT-expressing Ntsr1+ neurons in cortical layer 6, we used ambient amber light surrounding the mouse to mask sight of the optogenetic amber LED/laser.

### Data analysis: sorting single units

#### Multi-unit activity

Multi-unit activity was the high-pass-filtered electrophysiology signal (above 300 Hz).

#### Cluster selection

First, we clustered spike waveforms into clusters using UltraMegaSort (D.N. Hill, S.B. Mehta and D. Kleinfeld) by dividing each linear array of 16 electrodes into 4 tetrodes (or 32 electrodes into 8 tetrodes; 4 neighboring electrodes in each tetrode) and then clustering spikes based on the shape and size of spike waveforms on each tetrode. We selected well-isolated clusters with spike waveform amplitudes well above the spike detection threshold, which was set at 5.5 times the standard deviation of the high-frequency noise. We fit a Gaussian to the distribution of spike waveform amplitudes in each selected cluster and verified that more than 85% of the spikes in the cluster were above the spike detection threshold. Furthermore, we ensured that, within each cluster, there were fewer refractory period violations (i.e., cases when two spikes occurred within 1.5 ms of each other) than 1% of the number of total spikes.

#### Clustering bursts

Relay neurons in the dLGN sometimes burst. These bursts include a few spikes from the same neuron, very close together in time, and the amplitude of the spike waveform decreases for each spike in the burst, but the spike waveform shape does not change. Because UltraMegaSort uses spike amplitude to cluster the spikes, it is conceivable that smaller burst spikes from a dLGN unit might be clustered separately from the non-burst spikes and from the first spike in the burst. Our goal was to be sure that each single unit contained the burst as well as non-burst spikes from the neuron (for example, see Figure Supplement 5 B). Therefore we used the following procedure to combine clusters of burst and non-burst spikes likely belonging to the same source, a single neuron. After selecting well-isolated clusters (see above), we inspected all pairwise combinations of these clusters and permanently combined the two clusters, e.g., cluster 1 and cluster 2, if the following conditions were met:

1. The average waveforms of cluster 1 across the tetrode scaled very precisely, as determined by-eye, onto the average waveforms of cluster 2 (i.e., same shapes of spike waveforms, although spike waveform sizes may differ)
2. If one cluster in the pair had smaller-sized spike waveforms, this cluster contained fewer spikes than the cluster with larger spike waveforms (because smaller spikes should only occur after the first large spike in a burst)
3. More than 5% of the spikes in the cluster with the smaller spike waveforms should immediately follow (i.e., occur within 10 ms of) a spike in the cluster with larger spike waveforms
4. The combined cluster (composed of the cluster pair) contained fewer refractory period violations than 1% of the total number of spikes in the combined cluster.

We iterated this procedure on the clusters that emerged after a first round of combinations, and iterated again on the output of that round of combinations, and so forth, until all pairs of clusters meeting the above criteria had been combined and no more clusters could be combined according to these criteria. We inspected this final set of clusters by-eye and validated only the clusters with spike amplitude modes well above the spike detection threshold and with fewer refractory period violations than 1% of the total number of spikes in the cluster. These final, curated clusters were the single units used for further analysis.

#### Sorting cortical units

For extracellular units recorded in cortex, we sorted the spike waveform clusters as described above, but we did not further combine clusters to find the burst spikes.

#### Separating regular-spiking (RS) and fast-spiking (FS) units in cortex

We measured the half-width of a spike waveform at its half-maximum height. We separated the regular-spiking (putative excitatory) units and fast-spiking (putative inhibitory) units in the cortex as described in (Reinhold et al., 2015).

### Data analysis: Separating the alert and quiescent brain states

A 16-site or 32-site linear extracellular recording electrode was placed at a depth such that the top channel on this electrode would sit in the ventral part of hippocampus and the bottom channels on this electrode would penetrate dLGN (see “Extracellular electrophysiology” above). Thus, we could measure the local field potential (LFP) from the hippocampus and record single units in dLGN simultaneously. We measured the theta power in the hippocampal LFP to distinguish the alert and quiescent brain states. We used the mtspecgramc function in Chronux (Bokil et al., 2010) to measure the theta power of the hippocampal LFP. We defined the theta power as the power between 5 and 7 Hz minus the power at 2 and 4 Hz, as in (Bezdudnaya et al., 2006). We calculated the power in each of these frequency bands over 0.8 s-long time windows by passing the following parameters to Chronux: movingwin=[0.8 0.2] params.tapers=[3 5].

Finally, we normalized the power spectrum by its sum across all frequencies at each time point to focus the analysis on the relative power in different frequency bands. For each experiment, we plotted a histogram of the theta power averaged across trials in the experiment. Note that silencing cortex does not change the theta power in hippocampus (see Figure Supplement 4 A); thus, we included time windows when the blue LED was illuminating cortex. We inspected this bimodally distributed histogram of theta power (e.g., Figure 1 B, scatter plot) and selected, by-eye, the theta power threshold that seemed to best divide the two modes in the histogram. We classified trials with an average theta power above this threshold as “alert” and trials with an average theta power below this threshold as “quiescent”. The mice were often but not always running in the alert state. For example, in Figure 1, each trial lasted for 14.5 seconds. Of the trials classified as alert, 86% of these theta trials showed running. Here “running” means that the mouse ran continuously for at least 1 second. When calculating fraction of time spent in each brain state, we smoothed the theta power using 4 second bins.

### Data analysis: Measuring cortical synchronization

We recorded the LFP in V1 using an extracellular electrode (see above). We then calculated the cortical synchronization as the low-frequency LFP power (0.5 to 6 Hz) minus the high-frequency LFP power (30-80 Hz). We used the mtspecgramc function in Chronux to measure the power over 0.8 s-long time windows and passed this function the following parameters: movingwin=[0.8 0.2] params.tapers=[3 5]. We normalized the power spectrum at each time point by its sum across all frequencies.

### Data analysis: Units of hippocampus theta and cortex synchronization

In general, the units of the power spectrum of the LFP are V^2 Hz^-1. However, in the cases of hippocampus theta and cortex synchronization, we normalized the power spectrum by its sum across all frequencies at each time point, in order to focus on the relative power across different frequency bands. Therefore these are normalized measures.

### Data analysis: Quantifying power of the F1 response

#### F1 response to full-field flicker at varying temporal frequencies

We presented full-field visual flicker at frequencies between 1 and 60 Hz. The stimulus structure was the following: 1 s of mean-luminance baseline (no flicker), 2 s of full-field flicker modulated sinusoidally in time, then another 2 s of mean-luminance baseline. We calculated the unit’s trial-averaged peri-stimulus time histogram (PSTH), a binned count of spikes per second, as the unit’s response. We measured the F1 response to this full-field flicker using the mtspectrumpb function in Chronux (Bokil et al., 2010) over the full 2 s of the time-varying visual stimulus. We passed the following parameters to mtspectrumpb: params.tapers = [0.9 2 0]. We measured the F1 response as the power in the response frequency matching the stimulus temporal frequency. We averaged the PSTH of the unit’s spiking before taking the power.

#### F2 response

In Figure Supplement 2 F-G, we measured the F2 response as the power at the response frequency equivalent to 2 times the stimulus temporal frequency.

#### F1 response to drifting gratings

We measured the power of a single unit’s F1 response to a drifting grating visual stimulus with a fundamental temporal frequency of 3 Hz as follows. We calculated the unit’s trial-averaged PSTH, a binned count of spikes per second, as the unit’s response. We then used the mtspecgrampb function in Chronux (Bokil et al., 2010) to measure the power at 3 Hz in that unit’s response. We averaged the PSTH of the unit’s spiking before taking the power. Unless otherwise noted, we measured the power at 3 Hz over a time window of 1 s. We then stepped this 1 s-long time window along the PSTH, at intervals of 0.05 s, to get a smoothly varying F1 power over time. We passed the following parameters to Chronux: movingwin = [1 0.05] params.tapers = [5 6] (note that the first number, 5, is the time-bandwidth product). When we measured the response to a visual stimulus with a temporal frequency of 1 Hz, we used a 2 second time window for the power calculation (see below “F1 response to full-field flicker at varying temporal frequencies” for discussion of frequency response curves as in Figure 1 E). We present the F1 power as follows. In the figures, we show the raw F1 power of spiking activity in units of 1e5. An F1 power of 0.1 in units of 1e5 (i.e., 1e4) roughly corresponds to a PSTH with a 10 spikes per second peak-to-trough sinusoidal modulation at 3 Hz only; an F1 power of 4 in units of 1e5 corresponds roughly to a PSTH with a 20 spikes per second peak-to-trough sinusoidal modulation at 3 Hz; and an F1 power of 16 in units of 1e5 corresponds roughly to a PSTH with a 40 spikes per second peak-to-trough sinusoidal modulation at 3 Hz.

#### Power calculation before or after trial-averaging

Unless otherwise specified, we measured the F1 power of the trial-averaged PSTH. However, we also measured the F1 power of each individual trial before averaging across trials to control for any change in eye position across trials, although rodents make more, not fewer, eye movements when running (Wallace et al., 2013), and this should, if anything, lower F1 amplitude during the alert as compared to the quiescent state, whereas we see higher F1 amplitude during the alert state. Also note that this method is more likely to pick up spontaneous low-frequency activity as spurious 3 Hz visually evoked signal. All results were qualitatively unchanged using either analysis approach (not shown).

### Data analysis: Normalization of F1 power

In some cases, we present the F1 power across units normalized to a certain condition, for example, in Figure 1. First, we subtracted the baseline F1 power before the onset of the visual stimulus. Second, each unit’s F1 response was divided by its total power across all response frequencies between 0 and 250 Hz. Third, we normalized the response curve across all temporal or spatial frequencies to the average response within a certain condition (e.g., control condition). We always confirmed that effects shown in the normalized plot qualitatively matched the effects on the raw F1 power, and we always show the raw data as well as normalized data including Figure Supplements.

### Data analysis: Normalizing single-unit peri-stimulus time histogram (PSTH) of F1 power

Unless otherwise specified, the average PSTH of F1 power was presented as an average across all single units. Where specified as normalized, the PSTH of F1 power was divided by the average visually evoked F1 power (i.e., during the visual stimulus) in the control condition (no optogenetic manipulation), in order to show the fractional change as a result of the optogenetic manipulation. We normalized each single unit’s F1 power to its own visually evoked F1 power in control conditions after subtracting the baseline F1 power. Note that, in the case that a unit had a small visually evoked F1 response with respect to baseline, this normalization amplified noise. Thus, we chose to include only the most visually responsive units in the normalized PSTH. These were the units with the largest increase in F1 power with respect to baseline (see next section).

### Data analysis: Selecting visually responsive units

When we presented the raw, non-normalized data, we included all units recorded in dLGN. However, when we presented a normalized PSTH of F1 power, we selected only the visually responsive units, because including non-visually responsive units in the normalized figure amplifies noise. To pick the visually responsive units, we chose a threshold F1 power and included all units with an F1 response greater than this. We chose a threshold to separate the top 25% of units with the largest F1 responses. These were the units deemed “visually responsive”. When we show figures including only visually responsive units, we always also show, somewhere else, the same experiment including all units recorded in dLGN (even units without a visual response). Thus, we always show both the result including all units as well as the result including only the visually responsive units. This occasionally produced differences in the exact magnitude of the effect but no change in the qualitative effect.

### Data analysis: Measuring the power spectrum of the S1 LFP

We measured the LFP in somatosensory cortex (S1) using an extracellular electrode (see above). We calculated the power spectrum of the S1 LFP using the mtspectrumc function in Chronux.

We passed the following parameters to this function: params.tapers = [3 5]. The LFP amplitude is given in Volts. The plot in Figure Supplement 4 C shows the power times the frequency on the X axis squared, in order to make higher frequencies easier to see on the plot.

### Data analysis: Fraction burst spikes

To test whether dLGN units that burst more also showed a bigger effect of cortical silencing, we calculated the fraction of each unit’s spikes that occurred in bursts. For instance, a “fraction burst spikes” of 0.1 for a unit indicates that 10% of all its spikes occurred in bursts.

### Data analysis: Selecting putative Ntsr1+ single units

#### ArchT or GtACR2

We aimed to quantify the fraction of Ntsr1+ units in cortex suppressed by the ArchT or GtACR2 optogenetic manipulation. Because we injected Cre-dependent ArchT or Cre-dependent GtACR2 into layer 6 of the visual cortex of the Ntsr1-Cre line, we expected a fraction of Ntsr1+ neurons to express ArchT or GtACR2 and therefore be suppressed by illumination with amber/blue light. However, an alternate source of suppression of cortical neurons needed to be considered: our data showed that suppressing cortical feedback onto thalamus led to a decrease in thalamic spontaneous activity, which is expected to lead to a decrease in cortical spontaneous activity (Reinhold et al., 2015). Thus, cortical units suppressed by illumination with amber light were likely to include both directly suppressed ArchT- or GtACR2-expressing Ntsr1+ units and cortical units driven by thalamic input. To study this, we considered the ArchT experiments. To identify the directly suppressed ArchT-expressing Ntsr1+ units, we noted that these units should be suppressed before dLGN spontaneous activity was suppressed. We measured the time course of suppression of spontaneous activity in dLGN and found that its onset was delayed by 35 ms with respect to the onset of the amber LED directed at cortex (Figure Supplement 8 B,C). Hence we searched for units in the deep layers of cortex that were suppressed within the first 35 ms of the onset of the amber LED. These units represented putative ArchT-expressing Ntsr1+ neurons. (We used the same criterion for selecting GtACR2-expressing, putative Ntsr1+ units in visual cortex.)

For recordings in visual cortex in the Ntsr1-Cre mice, we positioned the 16-channel recording electrode in V1 such that the bottom-most channels on the electrode array were ventral to cortex (i.e., no spiking), and the electrode was placed at an angle of approximately 40 degrees from the tangent to the pia. We then measured putative layer 6 as the bottom-most 400 μm on the angled recording electrode, with respect to the deepest well-isolated single unit. Accounting for the electrode angle, this represents the bottom-most 250 μm of cortex, consistent with the size of layer 6 in the mouse.

Single units were classified as putative ArchT- or GtACR2-expressing if they met the following criteria:

1. Recorded in putative layer 6
2. Regular-spiking (see above)
3. Mean firing rate significantly suppressed within the first 35 ms of the onset of the amber/blue LED (p-value<0.05 according to Wilcoxon rank-sum)
4. Or mean firing rate suppressed (p-value<0.2 according to Wilcoxon rank-sum) within the first 35 ms of the onset of the amber/blue LED and firing rate still suppressed (including n.s. units) throughout visual stimulation (versus dLGN activity, which is not clearly suppressed during visual stimulation).

We found 25 out of 157 putative layer 6 regular-spiking units that met these criteria for the ArchT experiments. Because the Ntsr1+ neurons are about 60% of the total neurons in cortical layer 6 (Bortone et al., 2014), our results suggest that approximately 25% of the Ntsr1+ neurons expressed ArchT and were directly inhibited by illumination with amber light. (Note that, given these criteria, chance would predict that we see only 11.5% of putative layer 6 regular-spiking units that meet the criteria.) Importantly, the putative ArchT-expressing Ntsr1+ units were suppressed even before we observed any suppression of spontaneous activity in dLGN.

### Data analysis: Difference in baseline firing rate of dLGN neurons across mouse lines

The average spontaneous firing rate of dLGN neurons in the Ntsr1-Cre mouse line (in the alert brain state, average: 5.4 Hz, s.e.m.: 0.46 Hz, n=144 units) was lower than in the vGat-ChR mouse line (in alert brain state, average: 7.3 Hz, s.e.m.: 0.34 Hz, n=377 units). This was a significant difference quantified using the Wilcoxon rank-sum test (p-value=0.0029). Despite this, we observed the same effect of suppressing cortico-thalamic feedback on dLGN visually evoked activity in both mouse lines.

### Data analysis: Defining preferred and non-preferred spatial frequencies (Figure Supplement 9 C,D,E)

To test how cortico-thalamic feedback affects the representation of spatial frequency in dLGN, we presented the spatial frequencies 0.02, 0.03, 0.04 and 0.08 cycles per degree. We then identified the preferred and non-preferred spatial frequencies for each dLGN neuron by measuring the F1 power (visually evoked and baseline-subtracted, i.e., over the entire duration of the visual stimulus minus the F1 power in the 4 seconds preceding the visual stimulus) in response to each spatial frequency and ordered these responses from highest (preferred) to lowest (non-preferred). We selected one preferred spatial frequency and one non-preferred spatial frequency for each neuron. For plots during the quiescent state, we identified the preferred and non-preferred spatial frequencies based on the response in the alert brain state during cortical silencing. For plots during the alert brain state, we identified the preferred and non-preferred spatial frequencies based on the response in the quiescent brain state during cortical silencing. (Thus we avoid any issues of regression to the mean. For example, if we had chosen the preferred and non-preferred visual stimuli based on the responses in control, we would expect, by chance, the tuning to be less strong during cortical silencing, by definition, as a result of regression to the mean. However, our approach, using an independent measurement to select preferred and non-preferred visual stimuli, avoids this confound.) Therefore we can compare spatial frequency tuning with and without cortical silencing.

### Data analysis: Selectivity index and tuning ratio for spatial frequency tuning

We present two metrics capturing spatial frequency tuning in the dLGN, the tuning ratio and the selectivity index (Figure Supplement 9 D,E). We defined the tuning ratio as the average visually evoked and baseline-subtracted F1 power in response to the preferred spatial frequency divided by this average in response to the non-preferred spatial frequency. We defined the selectivity index as the difference of the preferred response and the non-preferred response divided by the sum of the preferred response and the non-preferred response. We ran the statistical tests on the selectivity index metric to avoid “divide by 0” issues, but we also present the tuning ratio for intuition about the effect size.

---

**Figure Supplement 1.**
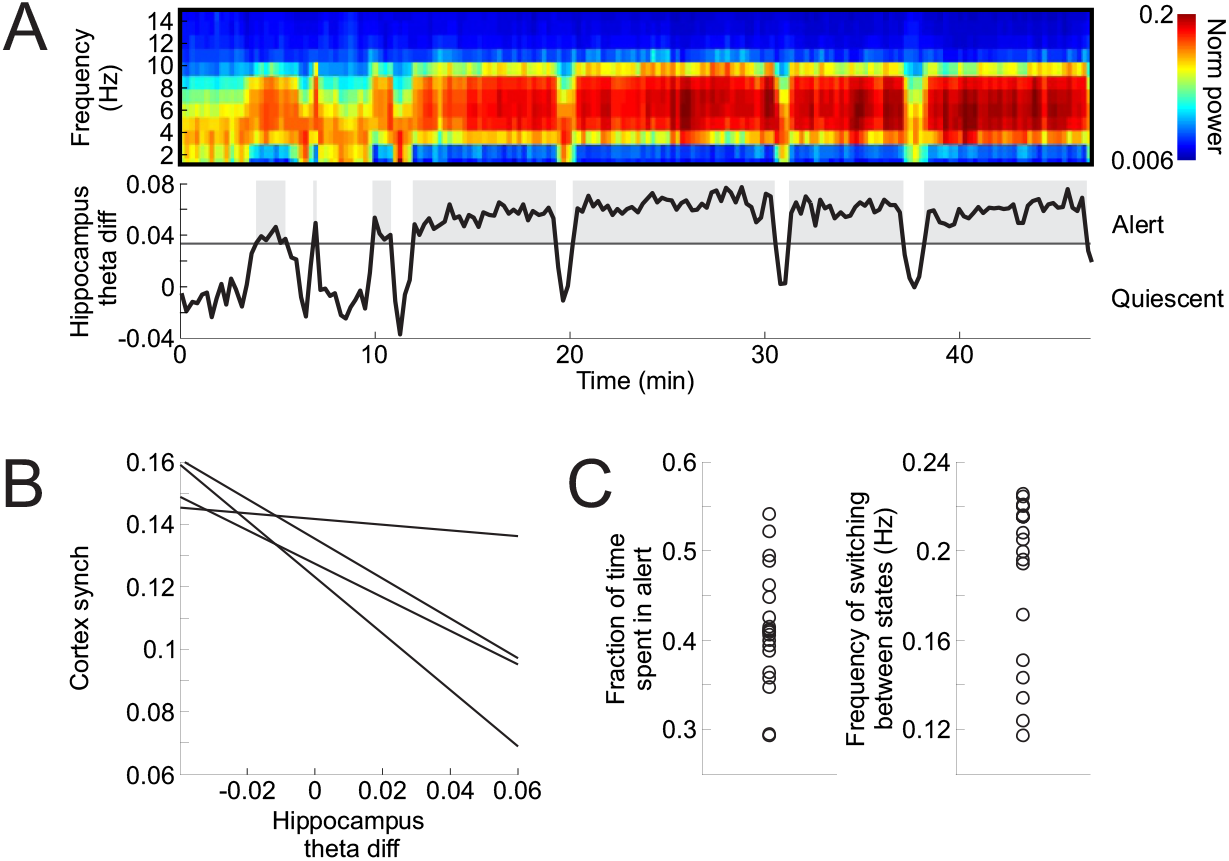
Definition of quiescent and alert brain states according to hippocampal local field potential (LFP). **(A)** Power spectrum of hippocampal LFP and theta band activity in one example recording session in awake mouse. (Top row) Color matrix: Power of hippocampal LFP across frequency and time divided by the total power at each time point. (Bottom row) Black trace is hippocampus theta difference (theta diff), i.e., power at 5-7 Hz minus power at 2-4 Hz. Black solid line shows threshold for this experiment. Quiescent brain state is below threshold. Theta or alert brain state is above threshold. Gray shading: Alert states. **(B)** Inverse correlation between hippocampus theta difference, defined in (A), and cortical synchronization, defined as the LFP power at 0.5-6 Hz minus power at 30-80 Hz in V1, from 4 example sessions of simultaneous hippocampus and V1 recordings in four different mice. Each line is the linear fit to the scatter of 1 s-long data points showing the simultaneously recorded hippocampus theta difference versus cortical synchronization (as the dotted line in Figure 1 B). **(C)** (Left) Fraction of total recording session spent in alert state. (Right) Frequency of switching between alert and quiescent states. Each point is one recording session (n=20 sessions from 9 mice).

**Figure Supplement 2.**
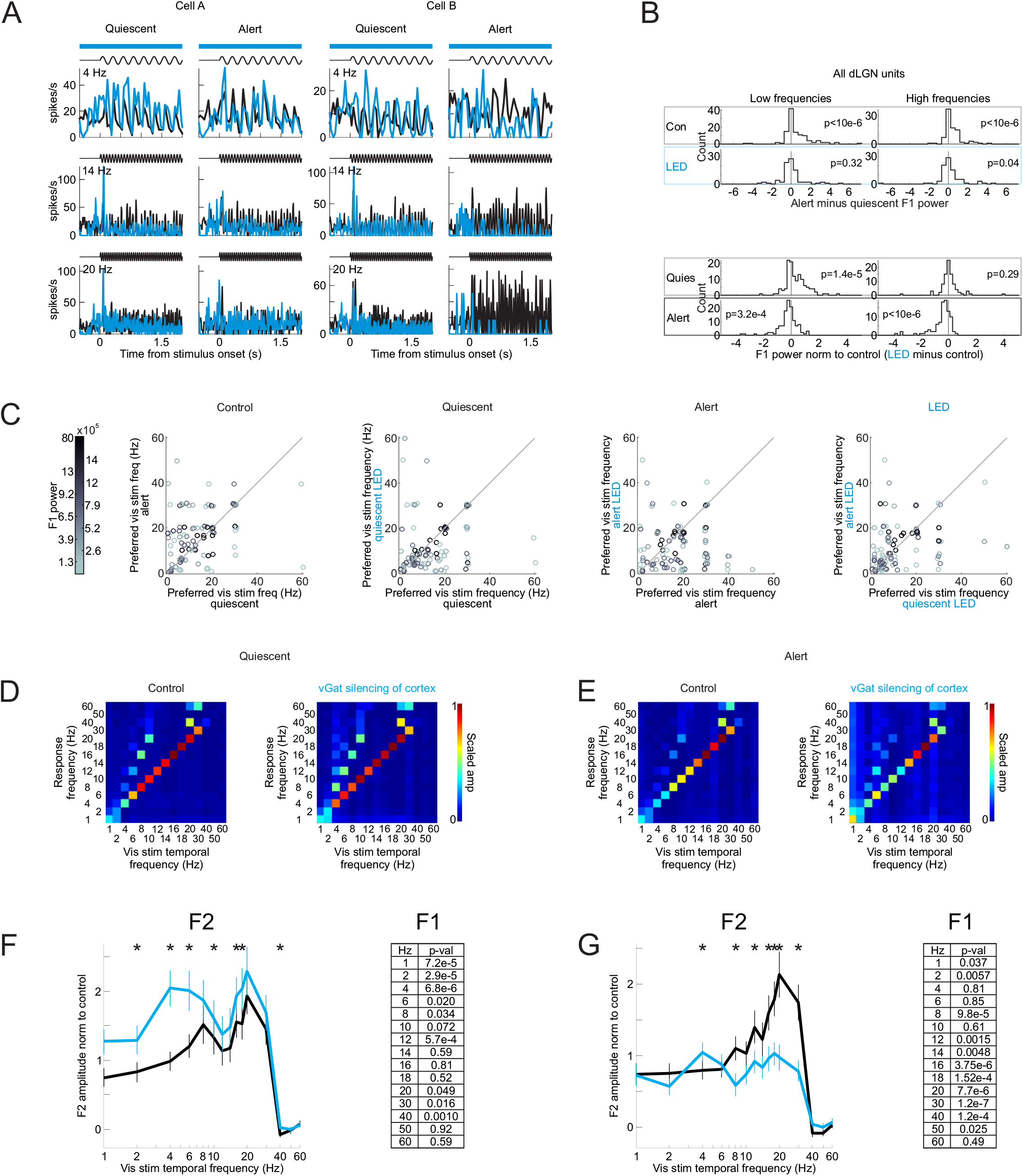
Silencing visual cortex and the effects on the dLGN frequency response across various temporal frequencies of the visual stimulus. **(A)** Trial-averaged visual responses (peri-stimulus time histogram, PSTH) of two example single units in dLGN. Setup as in Figure 1. Black: control. Blue: cortical silencing. Note that at low stimulus frequencies (4 Hz) cortical silencing increases the amplitude of the F1 response in the quiescent state, while at high stimulus frequencies (20 Hz) cortical silencing decreases the amplitude of the F1 response in the alert state. **(B)** Top rows: Distribution of alert minus quiescent F1 power for low (3 to 12 Hz; left) and high (13 to 40 Hz; right) stimulus frequencies in control (top) and cortical silencing (bottom). Distributions skewed toward positive values indicate that the alert F1 response was greater than the quiescent F1 response (n=254 from 5 mice). P-values are from Wilcoxon signed rank test. Note that with cortical silencing the distributions are centered around 0. Bottom rows: Distribution of cortical silencing minus control F1 power (normalized to control) for low (left) and high (right) stimulus frequencies in quiescent (top) and alert (bottom) conditions. Distribution skewed toward positive values indicates that the F1 response was larger during cortical silencing. Note the positively skewed distribution at low frequencies in the quiescent state and the negatively skewed distribution at high frequencies in the alert state. **(C)** Change in preferred frequency of dLGN units as a function of brain state and cortical silencing. Panel labeled “Control”: Preferred visual stimulus temporal frequency in quiescent versus alert (each dot is one unit). Panel labeled “Quiescent”: Preferred visual stimulus temporal frequency in quiescent state in control versus cortical silencing. Panel labeled “Alert”: Preferred visual stimulus temporal frequency in alert state in control versus cortical silencing. Panel labeled “LED”: Preferred visual stimulus temporal frequency during cortical silencing in quiescent versus alert (n=92 units from 3 mice). Color of dot is the power of that unit’s F1 response at the preferred visual stimulus frequency (color scale to the left). **(D)** Response of dLGN units to full-field flicker visual stimulus in the quiescent state. Average amplitude of the dLGN single-unit response to visual stimuli delivered at various temporal frequencies (n=92 units from 3 mice). Colors show amplitudes of the response scaled between minimal and maximal response (scale bar at right). (Left) dLGN response in control conditions. (Right) dLGN response during cortical silencing. Note F1 and F2 (i.e., response at a frequency double that of the visual stimulus) modulation of the response (diagonals in heatmap grids). **(E, G)** As in (D, F) but in the alert state. **(F)** Average and s.e.m. of amplitude of dLGN single-unit F2 response to full-field flicker stimuli in the quiescent state. Each unit’s response is normalized by the integral of that unit’s total response (sum of 15×15 heatmap grid in (D); Methods). Black: control. Blue: during cortical silencing. Asterisks indicate significant change with cortical silencing (p-values in table at right, paired Wilcoxon sign-rank test comparing control versus LED, n=92 units; asterisks indicate p<0.05, n=3 mice).

**Figure Supplement 3.**
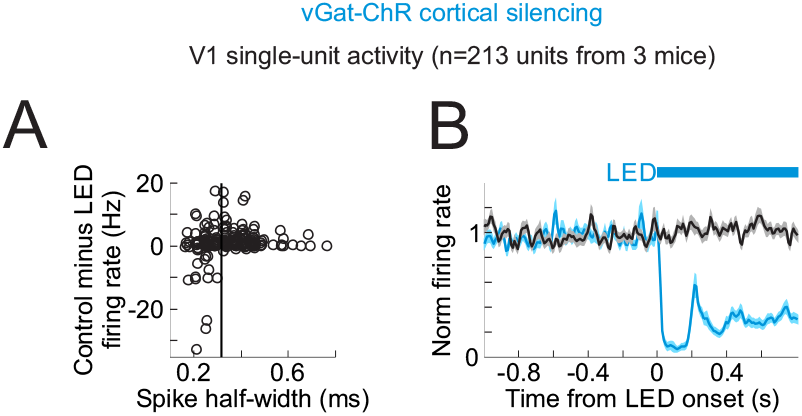
Efficiency of cortical silencing. **(A)** Impact of LED illumination on spontaneous activity of V1 units in vGat-ChR2 mice depending on spike shape (across all layers). Y axis is spontaneous firing rate in control minus firing rate during LED illumination of cortex. Negative values indicate increase in firing rate by LED. X axis is half-width at half-max of average spike waveform for each unit. Black vertical line indicates threshold for classifying units as regular-spiking or fast spiking (regular-spiking if spike half-width above threshold, n=213 units from 3 mice). **(B)** Time course of the impact of LED illumination on the spontaneous activity of regular spiking units in V1 of vGat-ChR2 mice. Average and s.e.m. from 213 units from 3 mice. Black trace is control; blue is with LED illumination. Blue bar indicates duration of LED illumination. All single unit firing rates normalized to 1 s baseline preceding LED onset.

**Figure Supplement 4.**
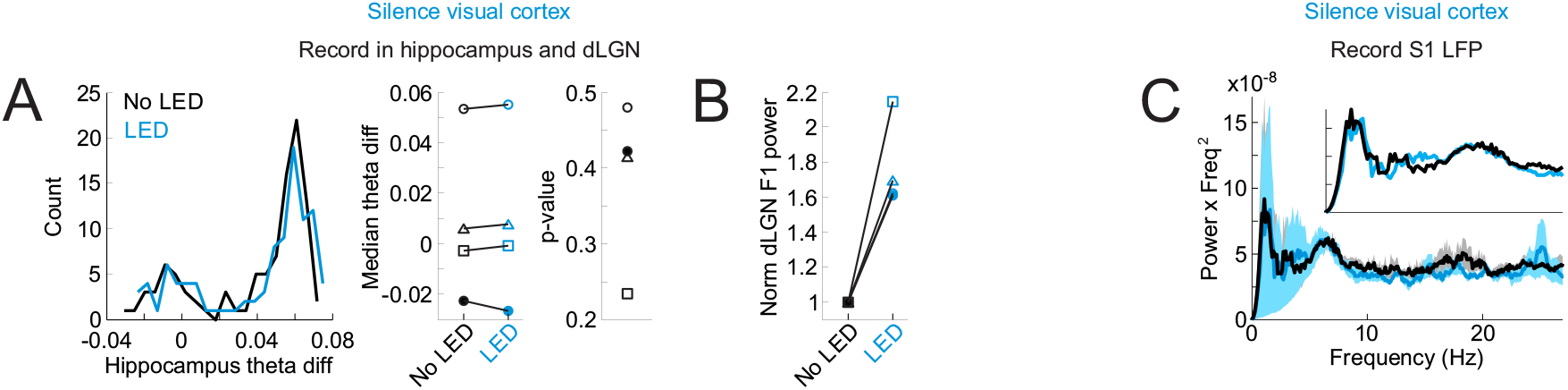
Silencing visual cortex does not alter brain state. **(A)** Left: Histogram of hippocampus theta difference (theta diff: power at 5-7 Hz minus power at 2-4 Hz) across 14.5 s-long intervals from an example mouse, with or without LED illumination of V1 to silence cortex. Center: Median hippocampus theta difference across a recording session during time periods with or without LED illumination of V1 to silence cortex. Each pair of points is one mouse. Right: P-values of Wilcoxon rank-sum comparison of hippocampus theta difference during time periods with or without LED. No change. **(B)** Effects of cortical silencing on visually evoked dLGN F1 power in the same 4 mice as in (A). Symbols as in (A). dLGN F1 power averaged across units and normalized to control. Note clear change in dLGN F1 power. **(C)** No change in local field potential (LFP) in somatosensory cortex (S1) during silencing of visual cortex. Average and s.e.m. of power spectrum of S1 LFP across 3 recordings from 2 mice. Power is shown here as power multiplied by the frequency (on the X axis) squared (whitening, Methods). Black is control. Blue is on interleaved trials during silencing of visual cortex including quiescent and alert trials. Inset is close-up between 1 and 10 Hz.

**Figure Supplement 5.**
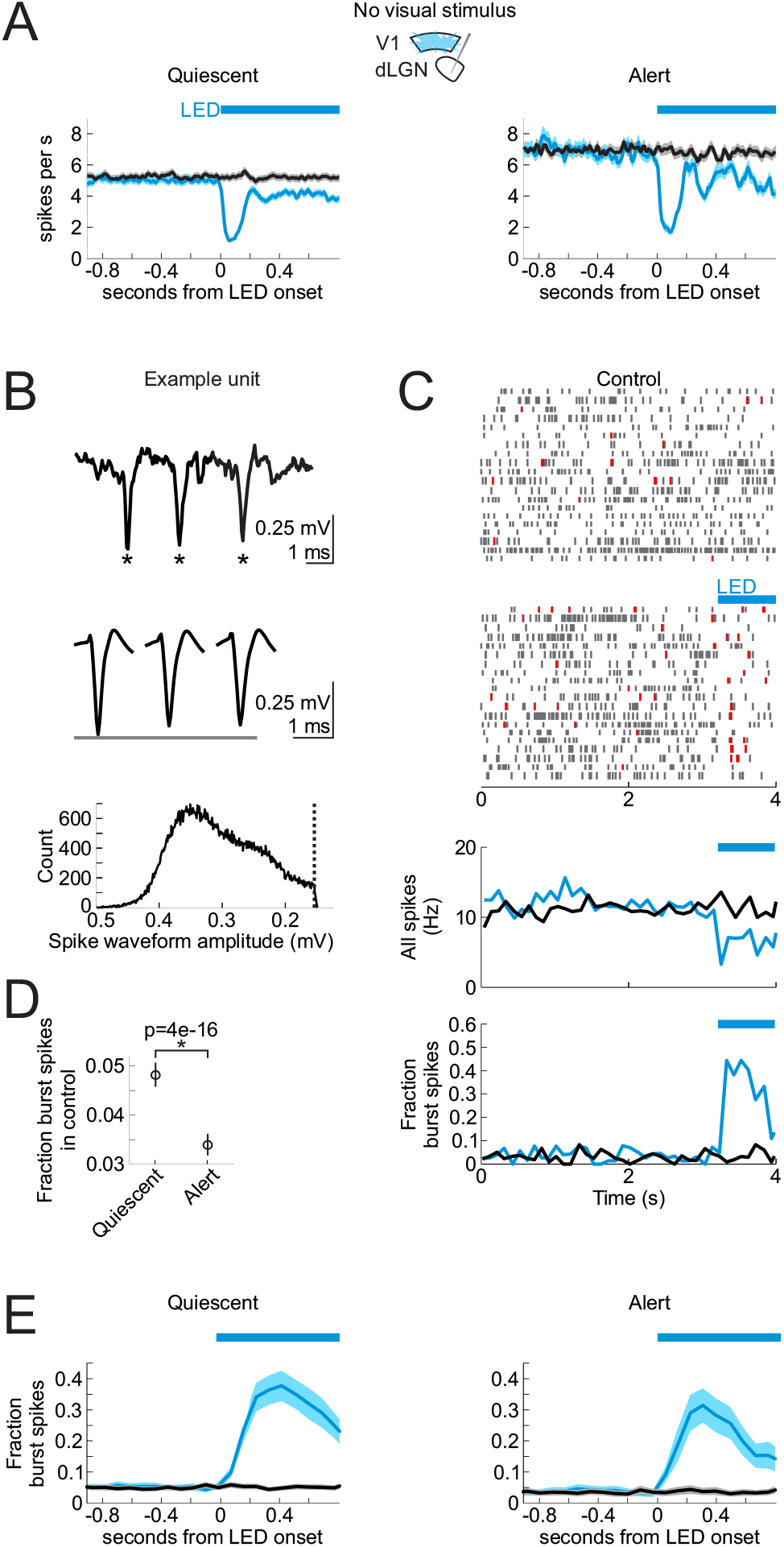
Silencing V1 decreases the spontaneous activity and increases bursting of dLGN neurons in alert and quiescent brain states. **(A)** Time course of the impact of V1 silencing on the spontaneous activity of dLGN units in the quiescent (left) and alert (right) state. Plots are average and s.e.m. (n=377 units from 6 mice). Black: control. Blue: during cortex silencing in vGat-ChR2 mice. Blue bar above plots indicates the duration of LED illumination of visual cortex. **(B)** Top: Example raw trace of spike burst from dLGN unit. Asterisks mark spikes in burst. Center: Average waveforms of first, second and third spikes in burst from this same dLGN unit. Bottom: Histogram of spike waveform amplitudes across all spikes from this unit. Dotted vertical line indicates spike detection threshold. **(C)** Response of example unit in (B) during quiescent state to onset of LED illumination of V1 to silence cortex (randomly selected example trials). Blue bar: LED illumination of V1 to silence cortex. Top: Spike raster plots without and with LED illumination. Red lines are bursts (Methods for details on detecting bursts). Bottom: Peri-stimulus time histograms of spiking in this dLGN unit, across all trials (including trials not shown in raster). Blue lines are trials with cortical silencing. Black lines are control. Bottom row shows the fraction of total spikes that occur in bursts for this same unit. **(D)** Average fraction of burst spikes during spontaneous activity (no visual stimulus) in quiescent versus alert brain states. P-value is from paired Wilcoxon sign-rank (n=377 units from 6 mice). **(E)** PSTH (average and s.e.m) of the fraction of all spikes that occur in bursts across all dLGN units (n=377 units from 6 mice) in control (black) and in response to cortical silencing (blue). Left: In quiescent state. Right: In alert state.

**Figure Supplement 6.**
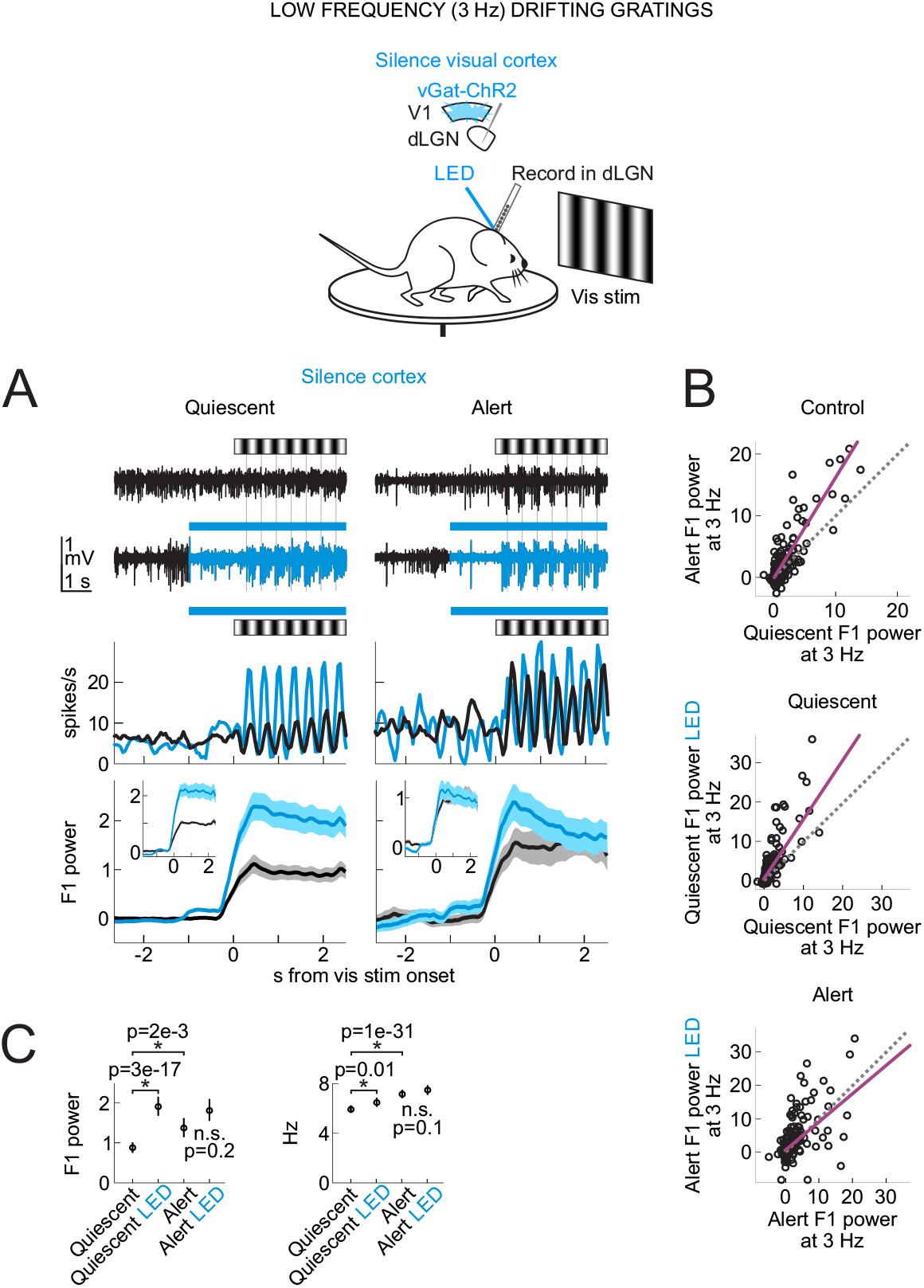
Silencing the visual cortex abolishes the brain state-specific differences in the visually evoked F1 response of dLGN neurons to drifting gratings. Schematic shows awake head-fixed mouse free to run or rest on a circular treadmill during electrophysiology recording, presentation of drifting gratings while silencing visual cortex by LED illumination of ChR2-expressing inhibitory cortical interneurons (vGat-ChR2). **(A)** Top: Example dLGN multi-unit (MU) traces before and during presentation of drifting gratings (0.03 cycles per degree at 3 Hz). Black: control. Blue: during cortex silencing. Middle: PSTH of example dLGN single unit. Bottom: Mean and s.e.m. of baseline-subtracted F1 power including all units (n=377 units from 6 mice, baseline is 4 s preceding visual stimulus). F1 power in units of 1e5 throughout this figure. Insets: mean and s.e.m. normalized to control F1 response including only visually responsive single units (Methods for selection of visually responsive units). **(B)** Baseline-subtracted F1 power of responses to drifting gratings. Each dot is a single dLGN unit. Purple line is linear fit to scatter. Dotted line is unity line. Top: Quiescent control vs. alert control. Middle: Quiescent control vs. quiescent with cortical silencing. Bottom: Alert control vs. alert with cortical silencing. **(C)** Average and s.e.m. across dLGN single units. Left: Baseline-subtracted F1 power. Right: Raw firing rates during visual stimulation (F0 response). P-values from paired Wilcoxon sign rank (n=377 units).

**Figure Supplement 7.**
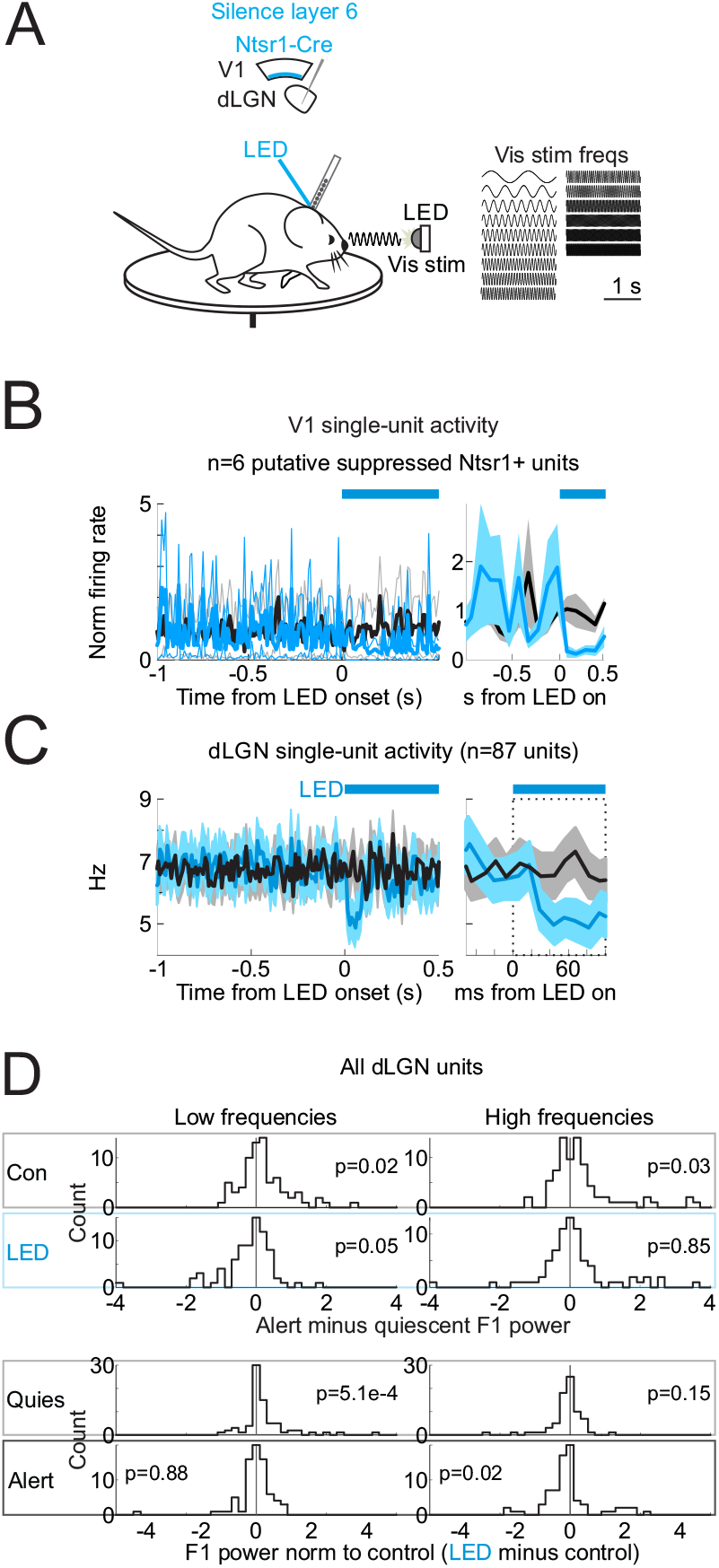
Silencing layer 6 cortico-thalamic neurons and the effects on the dLGN frequency response across various temporal frequencies of the visual stimulus. **(A)** Experimental set-up for silencing of layer 6 neurons expressing GtACR2. **(B)** Effects of LED illumination of visual cortex on spontaneous activity of putative Ntsr1+ cortico-thalamic feedback neurons in visual cortex (see Methods for definition of putative Ntsr1+ neurons). Left: Mean and s.e.m. firing rates of V1 putative Ntsr1+ neurons. Firing rates normalized to 1 s baseline preceding LED onset. n=6 units from 2 mice. Right: Close-up of left panel at LED onset. **(C)** Left: Effect of LED illumination of visual cortex on average and s.e.m. of spontaneous dLGN single-unit activity (n=87 units from 4 mice). Black: control. Blue: with illumination of cortex. Blue bar indicates duration of LED illumination of visual cortex. Right: Close-up of left panel at LED onset. **(D)** Layout showing effects on dLGN single units, as in Figure Supplement 2 B, except that here only layer 6 cortico-thalamic neurons in the visual cortex are suppressed using GtACR2.

**Figure Supplement 8.**
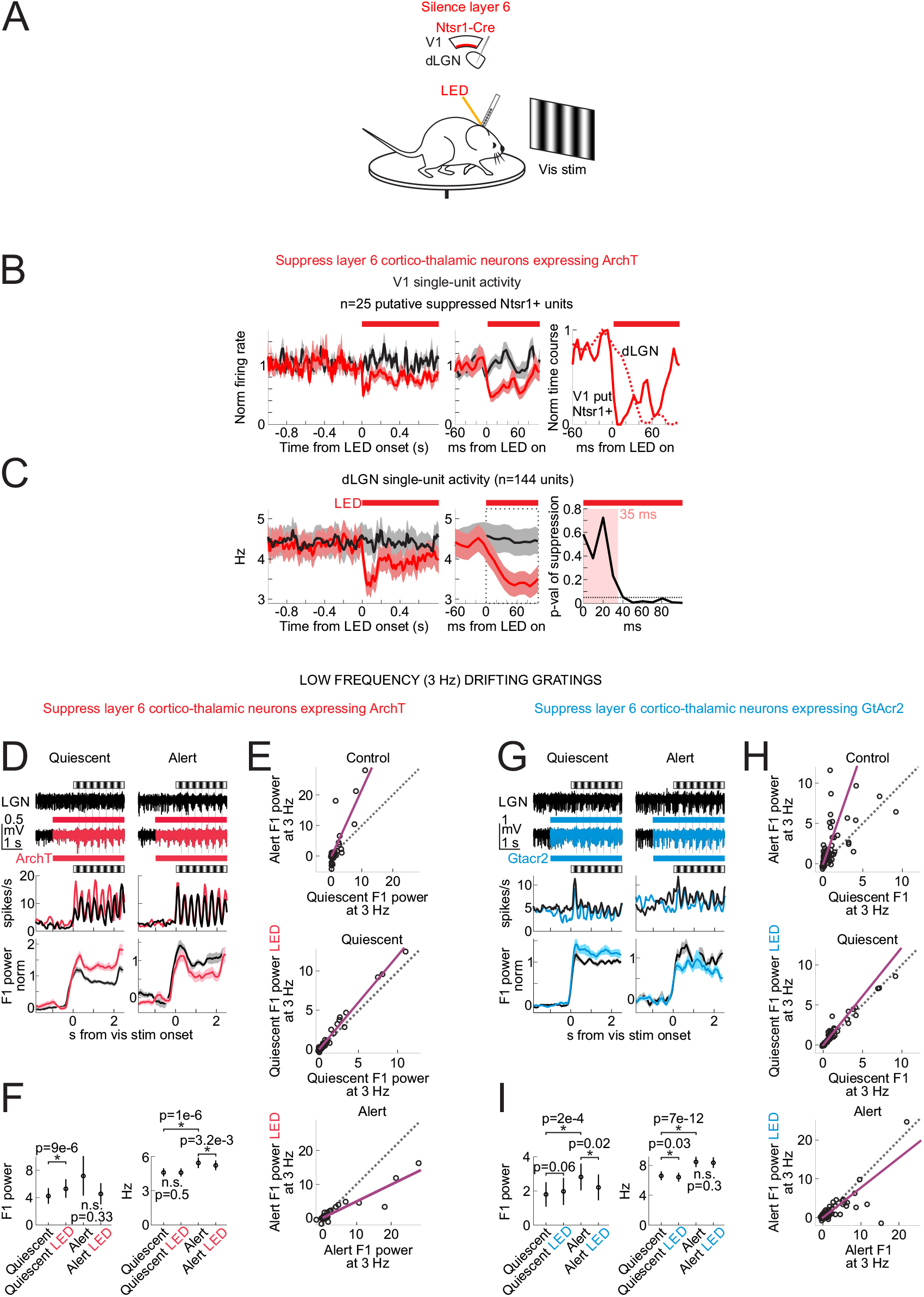
Silencing layer 6 cortico-thalamic neurons and the effects on the dLGN response to drifting gratings. **(A)** Experimental set-up for silencing of layer 6 neurons expressing ArchT. **(B)** Left: Mean and s.e.m. firing rates of V1 putative Ntsr1+ neurons. Firing rates normalized to 1 s baseline preceding LED onset. n=25 units from 4 mice. Middle: Close-up of left panel at LED onset. Right: Min-subtracted and max-normalized time course of change in average single-unit spontaneous activity. Dark red line is average of putative Ntsr1+ V1 units. Dotted red line is dLGN unit average from (B). Note fast suppression of putative Ntsr1+ V1 units preceding effects in dLGN. (See Methods for further discussion.) **(C)** Left: Effects of LED illumination of visual cortex on average and s.e.m. of spontaneous dLGN single-unit activity (n=144 units from 4 mice). Black: control. Red: with illumination of cortex. Red bar indicates duration of amber LED illumination of visual cortex. Middle: Close-up of left panel at LED onset. Right: P-value of paired Wilcoxon sign-rank test comparing firing rate in control (black) and during LED illumination of visual cortex (red) as a function of time from LED onset. Time window here refers to dotted square in middle panel. Significant change first observed after 35 ms post-LED onset. Dotted horizontal line is p=0.05. **(D-F)** Effects of layer 6 silencing on dLGN. Layout as in Figure Supplement 6 A,B,C but for selective suppression of layer 6 by ArchT. n=144 units from 4 mice. **(G-I)** As D-F but for selective suppression of layer 6 by GtACR2. n=87 units from 4 mice.

**Figure Supplement 9.**
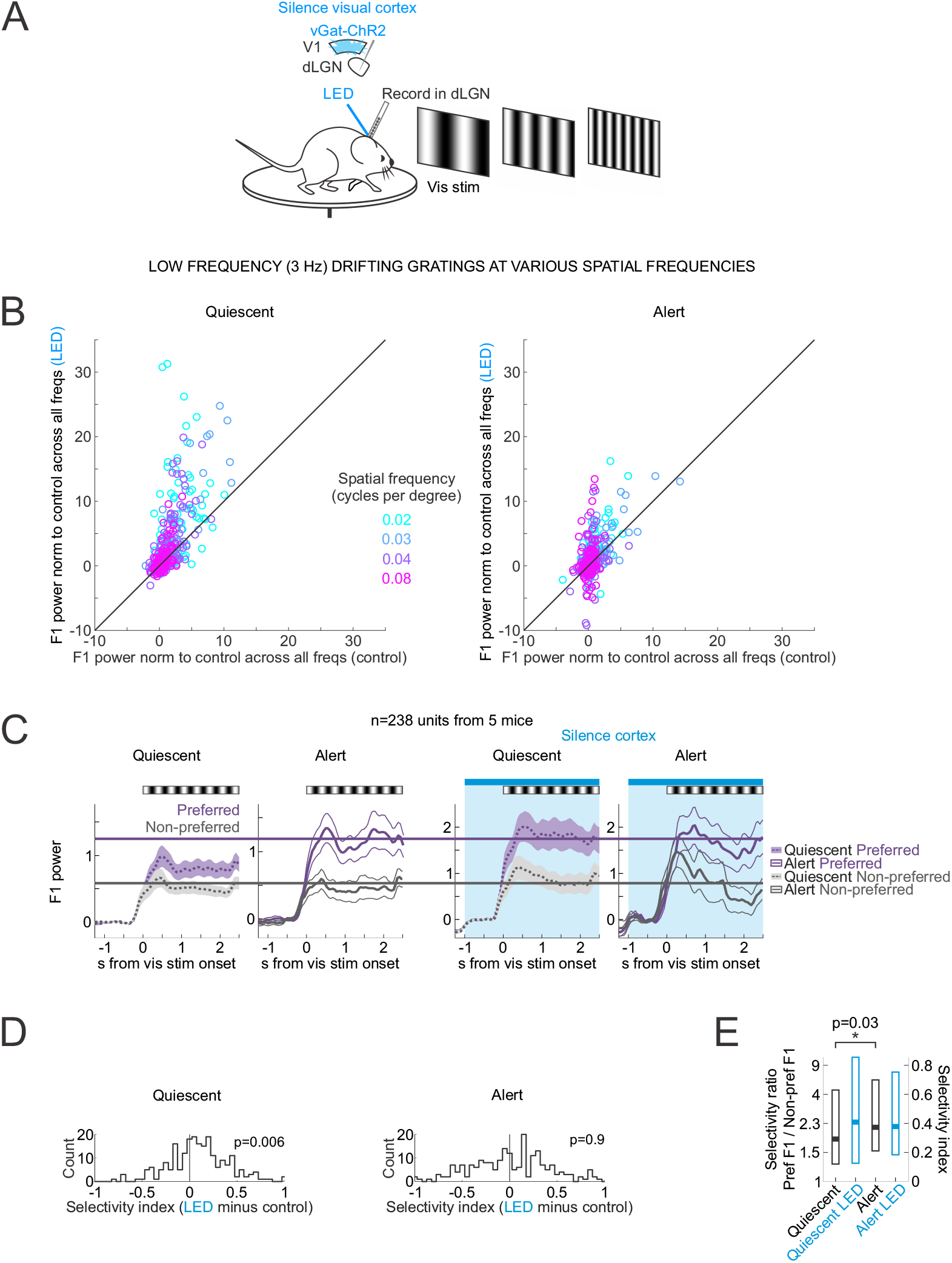
Silencing visual cortex abolishes brain state-specific differences in the spatial frequency tuning of dLGN neurons. **(A)** Experimental set-up to silence visual cortex while presenting drifting gratings at various spatial frequencies, as in Figure 3. **(B)** F1 responses of all dLGN units (n=238 units from 5 mice, each dot is one unit) in control versus cortical silencing (LED) at four spatial frequencies of the drifting grating (color coded). For each unit, F1 power was baseline-subtracted and normalized to control response (Methods). Left: Quiescent state. Right: Alert state. **(C)** Average and s.e.m. of power of F1 visual response to preferred (purple) or non-preferred (gray) spatial frequency across dLGN single units (Methods for definition of preferred/non-preferred). Responses are baseline-subtracted. F1 power in units of 1e5. Black and white stripes above plots show the duration of drifting grating. Blue bars show the duration of LED illumination of the visual cortex. Note selective enhancement of preferred response in the quiescent state upon cortical silencing. Purple and gray lines are for visual comparisons. **(D)** Effects of brain state and cortical silencing on spatial frequency tuning in dLGN. Histograms across all dLGN units (n=238). Selectivity index is the difference between the preferred F1 response and the non-preferred F1 response divided by the sum of the preferred F1 response and the non-preferred F1 response (Methods). Selectivity index is calculated for each unit. Shift of the distribution to the right indicates increased selectivity during cortical silencing. P-values are from paired Wilcoxon sign-rank test. **(E)** Summary of effects of brain state and cortical silencing on spatial frequency tuning in dLGN. Boxes show median with 25th and 75th percentiles. Selectivity ratio is the preferred F1 response divided by the non-preferred F1 response. Y axis on the right shows how the selectivity ratio maps onto the selectivity index (Methods). P-value from paired Wilcoxon sign-rank test.

**Figure Supplement 10.**
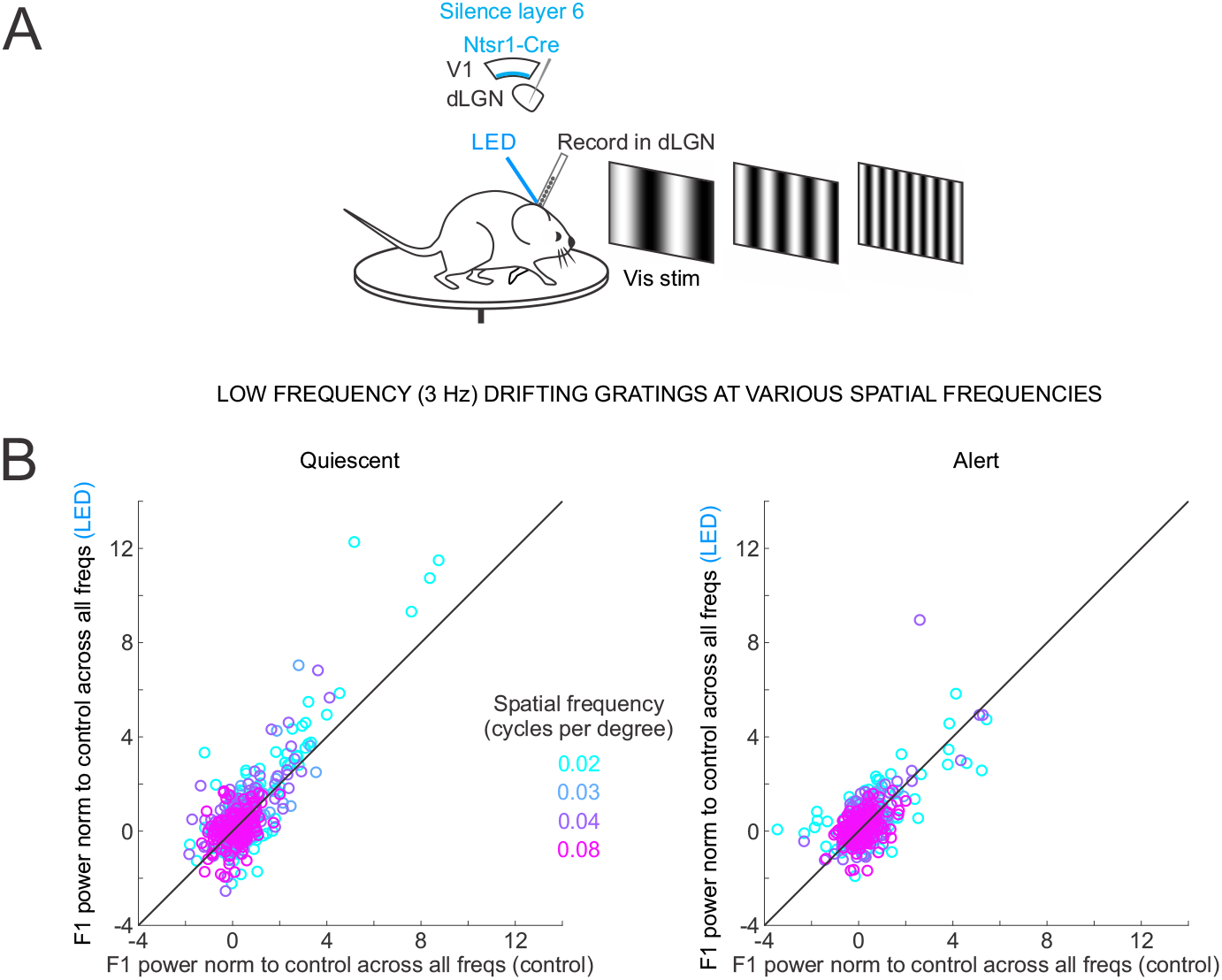
Silencing layer 6 cortico-thalamic neurons reduces brain state-specific differences in the spatial frequency tuning of dLGN neurons. **(A)** Experimental set-up to silence layer 6 cortico-thalamic neurons while presenting drifting gratings at various spatial frequencies, as in Figure 4. **(B)** F1 responses of all dLGN units (n=195 units from 5 mice, each dot is one unit) in control versus layer 6 silencing (LED) at four spatial frequencies of the drifting grating (color coded). For each unit, F1 power was baseline-subtracted and normalized to control response (Methods). Left: Quiescent state. Right: Alert state. Wilcoxon sign-rank comparing control to LED: 0.02 cpd in quiescent p-val=0.06, 0.03 cpd in quiescent p-val=0.4, 0.04 cpd in quiescent p-val=0.004, 0.08 cpd in quiescent p-val=0.3, 0.02 cpd in alert p-val=0.6, 0.03 cpd in alert p-val=0.8, 0.04 cpd in alert p-val=0.7, 0.08 cpd in alert p-val=0.8.

